# Cue overlap supports pre-retrieval selection in episodic memory: ERP evidence

**DOI:** 10.1101/2021.04.05.438462

**Authors:** Arianna Moccia, Alexa M. Morcom

## Abstract

People often want to recall only currently relevant events, but this selective remembering is not always possible. We contrasted two candidate mechanisms: the overlap between retrieval cues and stored memory traces, and the ease of recollection. In two preregistered experiments (*N*s = 28) we used event-related potentials (ERPs) to quantify pre-retrieval selection and the goal states – retrieval orientations – thought to achieve this selection. Participants viewed object pictures or heard object names, and one of these sources was designated as targets in each memory test. We manipulated cue overlap by probing memory with visual names (Experiment 1) or line drawings (Experiment 2). Results revealed that regardless of which source was targeted, the left parietal ERP effect indexing recollection was selective when test cues overlapped more with the targeted than non-targeted information, despite consistently better memory for pictures. ERPs for unstudied items were also more positive-going when cue overlap was high, suggesting that engagement of retrieval orientations reflected availability of external cues matching the targeted source. The data support the view that selection can act prior to recollection if there is sufficient overlap between retrieval cues and targeted versus competing memory traces.

## Introduction

We often want to retrieve a particular kind of event from memory. For example, we could check the reliability of a piece of information by recalling whether we heard it in conversation with friends or saw it on a news website. Ideally, we can selectively pull up relevant memories of news, without also recalling conversations with friends. Research suggests that people can only sometimes selectively remember in this way (Rosburg and Mecklinger, 2017). This ability tracks individual working memory capacity (Elward and Wilding, 2010) and is reduced in later life (Dywan, Segalowitz, and Webster, 1998; Keating et al., 2017). Achieving selective recollection depends on control processes that act prior to the point of memory retrieval. Time-resolved measures of brain activity like electroencephalographic event-related potentials (ERPs) allow us to quantify these proactive processes and their impact on recollection. Here, we investigated two factors proposed to be critical: the ease of retrieving targeted events (Herron and Rugg, 2003a), and the degree to which memory cues overlap stored information (Hornberger et al., 2004).

We used ERPs to quantify the retrieval of incidental as well as targeted information, and the goal states on which selection depends. In the recognition exclusion task (Jacoby, 1991) people study items in two sources, such as picture and word formats. They are then asked to focus retrieval on just one source at a time, accepting as targets only items from that source (e.g., those studied as pictures), and rejecting both items from the other source (non-targets, e.g. those studied as words) and unstudied (new) items. To do the task efficiently, participants need only to recollect items from the targeted source. Selective recollection is measured by comparing the left parietal ERP old/new effect for targets and non-targets (Dywan et al., 1998). This positive-going ERP modulation about 500-800 ms after the retrieval cue is well established as an index of recollection as opposed to familiarity in recognition tests (Rugg & Curran, 2007). When recollection is selective, the left parietal effect is larger for targets than non-targets, and non-target activity may be indistinguishable from new (for metanalysis see Rosburg and Mecklinger, 2017). This demonstrates that selection has occurred prior to retrieval. ERPs can also index goal-directed processes assumed to bring about selective remembering. When people adopt a retrieval orientation, they modify how retrieval cues are processed depending on the targeted source. We distinguished these goal states from successful retrieval by comparing ERPs for correctly rejected new items under different retrieval goals (Rugg & Wilding, 2000).

According to one view, selective recollection is only possible when target retrieval is easy (Herron & Rugg, 2003a). Several studies have demonstrated larger non-target left parietal effects when target accuracy was reduced, for example by manipulating study-test delay (Dzulkifli et al., 2006; Herron & Wilding, 2005), study list length (Wilding et al., 2005), or the encoding task (Herron & Rugg, 2003b). Relative target accuracy compared to non-targets may also be important: when non-targets are easier to recall, target recollection is typically not prioritized (Rosburg, Mecklinger, & Johansson, 2011). But accuracy does not seem to be the whole story. Larger left parietal effects to targets than non-targets are sometimes found even when target accuracy is low (Evans et al., 2010; Herron & Wilding, 2005; Sprondel, Kipp & Mecklinger, 2012).

An alternative proposal is that the ability to recollect selectively depends on how memory is cued. According to the encoding specificity principle, effective retrieval cues are ones that reinstate part of the information stored in the memory trace (Tulving and Thomson, 1973; see also Morris, Bransford, and Franks, 1977; Nairne, 2002). Hornberger et al. (2004) found positive-going retrieval orientation ERPs when people used visual word cues to retrieve words compared to pictures, and when they used picture cues to retrieve pictures compared to words. Thus, this goal-related brain activity tracked the overlap between test cues and studied items. Preliminary evidence suggests that cue overlap also has downstream consequences for recollection. Two studies found a target-selective left parietal ERP effect only when memory was probed with cues that exactly matched the target format, e.g., test cues and targets were visual words while non-targets were pictures (Herron & Rugg 2003a; Stenberg, Johansson, & Rosén 2006). However, these studies did not reveal whether the degree of overlap between cues and targets remains critical for selection when cues and targets are not identical. Clarifying this is critical because interactions between varied environmental cues and retrieval goals may impact people’s ability to draw efficiently on relevant memories in daily tasks. For example, selection from long-term memory contributes to working memory capacity (Unsworth, 2016). Effective external cues may also function as environmental support for memory, reducing the need for engagement of effortful proactive processes and benefiting populations with reduced executive function like healthy older adults (Craik, 1983; Morcom, 2016).

Here, we investigated whether cue-target overlap would enable selective recollection even when performance was consistently better for targets from one source, regardless of which was targeted. In two preregistered ERP experiments, participants had to remember either words they had heard or pictures they had seen, in separate blocks. In Experiment 1, test cues were visual words, which overlapped more with the auditory word source. Supporting the overlap view, the left parietal effect was larger for targets than non-targets only when auditory words were targeted. In Experiment 2, we used object line drawings as cues, which overlapped more with the picture source (Czernochowski et al., 2005). The findings confirmed a complementary asymmetry to Experiment 1, with greater selectivity again for the high-overlap source. In both experiments performance was better for the picture source, so the results cannot be explained by the ease of target recollection. As expected, the direction of ERP retrieval orientation effects also reflected the degree of cue-target overlap. Together these findings show that cue overlap enables selection prior to recollection, as predicted by the encoding specificity principle.

## Methods

### Participants

Twenty-eight participants were included in Experiment 1 (20 female, age *M* = 22.79 years, *SD* = 4.14), and other 28 in Experiment 2 (20 females, age *M* = 24.57 years, *SD* = 3.71). One further participant in Experiment 2 was excluded due to an insufficient number of artefact-free trials (for preregistered criteria see https://osf.io/j84z6 and https://osf.io/pqn4z/). Sample sizes were determined *a priori* using effect sizes from Dzulkifli and Wilding (2005). Power analysis using G*Power 3.1.9.2 indicated that 29 and 17 participants would be required respectively to replicate the smallest main effect of retrieval orientation, from 500-600ms (*d* = 1.4) and the main effect of target versus non-target left parietal effects from 500-800ms (*d* = 1.9) with .95 power at *α* = .05. *N* was rounded to 28 to simplify counterbalancing. Participants were right-handed with normal or corrected to normal vision and hearing, who were in good self-reported health and not taking medication that might affect cognition. All were very fluent in English (self-rated scores ≥ 15/20 on ratings adapted from Vega-Mendoza et al. (2015) (see https://osf.io/gcrm2/). The experiments were approved by the Psychology Research Ethics Committee at the University of Edinburgh, ref.: 135-1819/1 and 300-1819/1.

### Materials

Stimuli in both experiments were pictures and names of 240 common objects (Fig. 1). Study phase stimuli appeared as either colored pictures or as auditory words spoken by an English native male voice. At test, memory probes were visual words in Experiment 1, or grey-scale line drawings in Experiment 2. The audio files were a subset of those used by Hornberger et al. (2004). Corresponding object images were sourced from the BOSS database (Brodeur et al., 2014), POPORO database (Kovalenko et al., 2012), or online (see Supplemental Material available online). The critical items were divided into 6 sets of 40 items each. For each of the two study-test cycles, one set of pictures and one of auditory words were combined to create a study list of 80 items. The corresponding visual words (Experiment 1) or line drawings (Experiment 2) were then combined with a third set of new items to create the test list of 120 items. For each study-test cycle, half of the studied pictures, half of the studied auditory words, and half of the new items were allocated to the first test block, and the remainder to the second test block. In total, there were 80 critical targets, 80 critical non-targets, and 80 critical new items. Two further filler pictures were added at the beginning of each study list, and two unstudied filler items at the beginning of each test block. A further 12 items served in practice lists. Item presentation order was determined randomly within each study and test list.

**Fig. 1.**
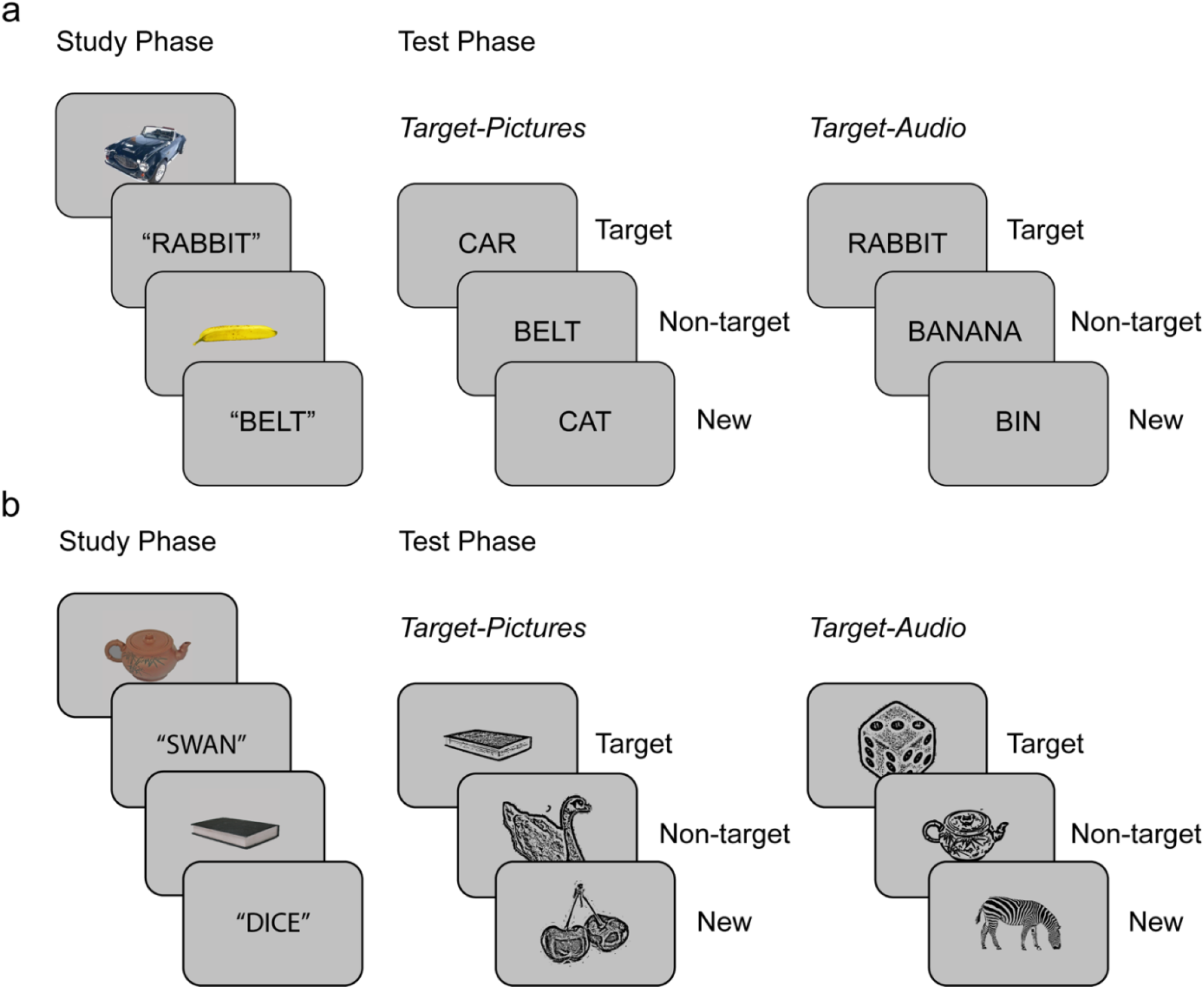
Experimental paradigm: recognition exclusion task procedure for Experiments 1 (a; visual word test cues) and Experiment 2 (b; line drawing test cues). In both experiments, participants studied a single block of pictures and auditory words. In each of the two test blocks, either pictures or auditory studied words were targets. Required responses were “yes” to the target items and “no” to non-target and new items (see Materials and Procedure for details).

### Procedure

Each experiment consisted of two study-test cycles (Fig 1), during which the EEG was recorded.

#### Study phase

Participants studied items presented as pictures or auditory words. Pictures appeared at the center of a square frame on a grey background subtending a visual angle of 4.32°. Auditory words were played at 44100 Hz, while a blank screen was shown. On each trial, a preparatory cue signaled the format of the upcoming item, either a yellow asterisk ‘*’ or a blue lowercase ‘o’ (allocation to pictures and auditory words was counterbalanced). Participants were instructed to learn the items for a subsequent memory test, while judging their pleasantness: “very pleasant”, “somewhat pleasant”, “pleasant”, or “not pleasant”. To maximize differences in processing between stimulus formats, they were instructed to take into account the holistic experience, paying attention to the visual or acoustic features. The preparatory cues were on-screen for 1,000 ms, followed by a blank screen for 100 ms, then stimulus presentation for 1,000 ms. A red fixation cross followed for 1,500 ms before the word “RESPOND” was presented at the center for up to 3,000 ms, during which participants were asked to give their answer. A 100 ms blank screen separated participants’ response and the next trial.

#### Test phase

Each test phase comprised two blocks with different target designations. In each, items presented in one format at encoding were designated as targets (Target-Pictures or Target-Audio). For example, in the Target-Pictures block, participants were instructed to answer ‘yes’ to an item if they had seen a picture of a corresponding object in the preceding study phase and ‘no’ to all other items. Target designation switched for the second test block and was also signaled on each trial using the same preparatory symbols as at study. All items appeared in the middle of the computer screen. In Experiment 1, test probes were visual words shown in 48 pt black uppercase letters. In Experiment 2, test probes were grey-scale line drawings, presented with a 3.71° visual angle. Test trials began with pre-cues for 500 ms, followed with a black fixation for 1,800 ms before stimulus presentation. Stimuli appeared for 3,000 ms followed by a red fixation for 500 ms in Experiment 1, and for 500 ms followed by a 3,000 ms fixation in Experiment 2. Participants’ responses were recorded during stimulus presentation in Experiment 1, and during fixation in Experiment 2. Participants were instructed to fixate in the middle of the screen throughout stimulus presentation, even after a response was made, to avoid excessive ocular movements.

The order of Target-Picture and Target-Audio blocks was counterbalanced across participants. Keypress responses used middle and index fingers at study, and index fingers at test, and the allocation of judgments to left and right hands was counterbalanced. The main experiment was preceded by a short practice phase, and study and test phases were separated by a 1-5 min interval, during which participants completed a distractor task consisting of 12 pen-and-paper true or false questions.

### EEG recording and pre-processing

EEG data were recorded with a BioSemi Active Two AD-box with 24-bit signal digitization from 64 active silver/silver chloride electrodes embedded in an elastic cap using the extended International 10-20 system configuration (Nuwer et al., 1998; http://www.biosemi.com/products.htm). Common Mode Sense and Driven Right Leg electrodes worked as ground electrode and noise rejection conjointly, while bipolar electrodes, placed above and below the right eye and on the outer canthi, recorded vertical and horizontal eye movements (electrooculogram; EOG). EEG and EOG signals were acquired continuously at a 1024-Hz sampling rate with amplifier bandwidth of 0±208 Hz (3 dB) and referenced to the CMS reference electrode. EEG data were preprocessed using the EEGLAB toolbox (Delorme & Makeig, 2004) in MATLAB R2018a. Data were first re-referenced offline to the average of the left and right mastoid electrodes. A 0.1-40 Hz Hamming windowed-sinc FIR filter was applied, with a 50 Hz notch filter for line noise. Data were divided into 4,500ms study and 6,700ms test epochs, time-locked to the stimulus onset. Customized threshold functions from the FASTER toolbox (Nolan et al., 2010) were used to identify and reject epochs and channels with excessive gross artefacts. The preprocessing pipeline and threshold criteria were developed for Experiment 1 and pre-registered for Experiment 2. Criteria were based on participant-level *z*-transformed values over trials that exceeded ± 3 (see Supplemental EEG preprocessing online for details). Independent Component Analysis (ICA) was used to correct for EOG artefacts, by manually removing ICA components attributable to vertical and horizontal eye movements. Rejected channels were then replaced by interpolation using data from neighboring electrodes. A 200 ms pre-stimulus baseline was used for ERP computation.

### Statistical analysis

All reported behavioral and ERP analyses were preregistered. Statistical analyses were conducted in R 3.6.1 (R Core Team, 2019) except where stated and alpha was set at .05. In analyses of variance (ANOVAs) we applied a Greenhouse-Geisser non-sphericity correction where appropriate. Benjamini-Hochberg false discovery rate multiple comparison corrections (Benjamini & Hochberg, 1995) were used in *post hoc* tests following significant interactions and all reported *p* values are adjusted. Cohen’s *d* was calculated by dividing mean differences by the pooled standard deviation (Dunlap et al., 1996).

## Results

### Exclusion task performance

We assessed differences in performance according to target designation for targets (items studied as pictures in the Target-Pictures condition or as auditory words in the Target-Audio condition), non-targets and new items.

**Table 1.** Recognition Exclusion Task Performance.

| Target Designation | Experiment 1 |  |  | Experiment 2 |  |  |
| --- | --- | --- | --- | --- | --- | --- |
|  | Target | Non-target | New | Target | Non-target | New |
| Target-Audio |  |  |  |  |  |  |
| Accuracy | .78 | .90 | .87 | .71 | .94 | .81 |
|  | [.74, .81] | [.88, .93] | [.84, .91] | [.67, .76] | [.92, .96] | [0.76, .86] |
| RTs | 1200 | 1257 | 1264 | 1315 | 1188 | 1529 |
|  | [1139, 1261] | [1222, 1293] | [1199, 1,330] | [1263, 1367] | [1136, 1240] | [1467, 1592] |
| Target-Picture |  |  |  |  |  |  |
| Accuracy | .81 | .86 | .92 | .89 | .88 | .91 |
|  | [.78, .85] | [.83, .90] | [.89, .94] | [.86, .91] | [.85, .91] | [.89, .93] |
| RTs | 1097 | 1308 | 1294 | 1062 | 1297 | 1310 |
|  | [1028, 1165] | [1249, 1367] | [1224, 1363] | [1008, 1116] | [1259, 1335] | [1234, 1385] |

Note: Table shows means and [95% confidence intervals] of accuracy proportions and median response times (RTs in ms) for correctly identified items according to trial type, target designation and experiment. Confidence intervals were adjusted using the Cousineau-Morey method for within-subject variables (Morey, 2008).

#### Accuracy

In Experiment 1, when retrieval cues were words, responses were generally more accurate for studied pictures, whether these were identified as targets (Target-Pictures) or non-targets (Target-Audio). ANOVA on accuracy proportions with factors of Item Type (targets/non-targets/new) and Target Designation (picture/audio) revealed a significant main effect of Item Type, *F*(1.59, 42.83) = 17.66, *p* < .001, *η*_*p*_^*2*^ = .395, a non-significant main effect of Target Designation, *F*(1, 27) = 0.90, *p* = .352, *η*_*p*_^*2*^ = 0.032, and a significant interaction, *F*(1.86, 50.34) = 5.35, *p* = .008, *η*_*p*_^*2*^ = 0.165. Although pairwise *post hoc t*-tests did not show significant effects of target designation in any condition, there was a slight accuracy advantage for items studied as pictures both for targets, *t*(27) = 1.91, *p* = .067, *d* = 0.40 and non-targets, *t*(27) = − 1.84, *p* = .076, *d* = 0.49, and for new items when targeting the pictures, *t*(27) = 1.79, *p* = .084, *d* = 0.48.

In Experiment 2, when retrieval cues were line drawings, accuracy was again greater for items studied as pictures. ANOVA with the same factors revealed significant main effects of Item Type, *F*(1.66, 44.78) = 14.67, *p* < .001, *η*_*p*_^*2*^ = 0.352 and Target Designation, *F*(1, 27) = 33.95, *p* < .001, *η*_*p*_^*2*^= 0.557, and a significant interaction *F*(1.81, 48.89) = 30.31, *p* < .001, *η*_*p*_^*2*^= 0.529. *Post hoc t*-tests confirmed more accurate identification of both targets and non-targets when they were studied as pictures than as auditory words, *t*(27) = 7.86, *p* < .001, *d* = 1.71, and *t*(27) = − 3.75, *p* = .001, *d* = 0.88. New items were also better identified when participants were targeting pictures, *t*(27) = 3.93, *p* = .001, *d* = 0.97.

#### Response times

We expected that participants would respond more slowly to non-targets than targets irrespective of targeted format, a pattern thought to suggest prioritization of target retrieval (Rosburg and Mecklinger 2017). We analyzed median RTs for trials attracting correct responses. When cues were words in Experiment 1, ANOVA with factors of Item Type (target hits/non-target correct rejections (CRs)/new CRs) and Target Designation (picture/audio) revealed a significant main effect of Item Type, *F*(1.47, 39.82) = 10.06, *p* = .001, *η*_*p*_^*2*^ = .272, a non-significant main effect of Target Designation, *F*(1, 27) = 0.08, *p* = .774, *η*_*p*_^*2*^ = .003, and a significant interaction, *F*(1.55, 41.97) = 7.41, *p* = .001, *η*_*p*_^*2*^ = .215. *Post hoc t-tests* confirmed that target responses were faster when pictures as opposed to auditory words were correctly identified, *t*(27) = − 2.6, *p* = .015, *d* = 0.61, while RTs for non-targets and new items did not differ significantly by target designation, *t*(27) = 1.45, *p* = 0.158, *d* = 0.41, and *t*(27) = 0.79, *p* = .434, *d* = 0.18.

Responses were also faster for items studied as pictures than auditory words in Experiment 2. ANOVA with the same factors revealed significant main effects of Item Type *F*(1.32, 35.72) = 28.80, *p* < .001 *η*_*p*_^*2*^ = 0.516, and Target Designation *F*(1, 27) = 27.07, *p* < .001, *η*_*p*_^*2*^ = 0.501, as well as a significant interaction *F*(1.92, 51.79) = 40.40, *p* < .001, *η*_*p*_^*2*^ = 0.599. *Post hoc t-*tests showed that participants were significantly faster when identifying items studied as pictures than auditory words, whether these were targets or non-targets, *t*(27) = − 9.03, *p* < .001, *d* = 1.85 and *t*(27) = 3.21, *p* = .003, *d* = 0.93. As for accuracy, responses to new items were also significantly faster when targets were pictures vs. auditory words, *t*(27) = − 5.39, *p* < .001, *d* = 1.60. Thus, participants recognized items they had studied as pictures faster than those studied as auditory words, despite the higher cue-target overlap with the auditory source in Experiment 1.

### ERP results

To test our principal hypotheses about the selectivity of target over non-target recollection, we examined the left parietal old/new effect in a focal analysis restricted to data from 3 parietal electrodes (P1/P3/P5) from 500 to 800 ms post-stimulus, following Dzulkifli and Wilding (2005). We quantified the mean per-participant stimulus-locked ERP amplitudes for correct responses in each experimental condition (target hits, non-target CRs, and new CRs) according to target designation (Target-Pictures and Target-Audio). These analyses were complemented by subsidiary, global analyses which tested whether ERPs evoked by targets and non-targets differed outside the predefined locations and time-windows. The global analyses included all electrodes and timepoints from 300-1,400 ms post stimulus, with a family-wise error correction using the nonparametric cluster permutation method from the FieldTrip toolbox (Maris & Oostenveld, 2007; Oostenveld et al., 2011). Thus, we i) ran dependent *t*-tests on the contrast of interest at each electrode and time-point, ii) defined clusters of temporally and spatially adjacent samples significant at *α* = .05 (sample-level *α*), iii) computed a cluster-level statistic equal to the sum of *t*-values per cluster, and iv) evaluated the maximum difference of this cluster-level statistic under its permutation distribution, created by randomly swapping data points between conditions within participants. We used 5,000 randomization draws to estimate each *p*-value (1-tailed cluster-level *α* of .025).

To assess retrieval orientation effects reflecting retrieval goal-states, we compared ERPs elicited by new CRs according to target designation. The focal analyses used a 3 x 3 grid of electrode locations (F5, Fz, F6/C5, Cz, C6/P5, Pz, P6) in three epochs: 300-600 ms, 600-900 ms, and 900-1200 ms, following Hornberger et al. (2004). Global analyses were also conducted following the above procedure. Additional analyses of retrieval orientation effects time-locked to the preparatory cues did not yield significant results and are included in Supplemental Results available online.

### Recollection selectivity

#### Focal analyses

When retrieval cues were words (Experiment 1), left parietal old/new effects for targets were larger than those for non-targets only in the high cue-target overlap condition: when targets were auditory words (Fig. 2a, c). ANOVA with factors of Item Type (target hits/non-target CRs/new CRs) and Target Designation (picture/audio) showed significant main effects of Item Type *F*(1.91, 51.60) = 24.64, *p* < .001, *η*_*p*_^*2*^ = .477, and Target Designation *F*(1, 27) = 20.10, *p* = .001, *η*_*p*_^*2*^ = .427, as well as a significant interaction, *F*(1.83, 49.40) = 3.32, *p* = .048, *η*_*p*_^*2*^ = .110. *Post hoc t-*tests for the Target-Pictures block showed that ERPs evoked by both target hits and non-target CRs were significantly more positive-going than those for new CRs, *t*(27) = 4.77, *p* < .001, *d* = 0.64, and *t*(27) = 3.36, *p* = .004, *d* = 0.51, while target and non-target ERPs did not differ significantly, *t*(27) = 0.29, *p* = .775, *d* = 0.04. Thus, recollection as measured by the left parietal effect was not selective for targeted information. In sharp contrast, in the Target-Audio block, ERPs for target hits were significantly more positive than for both non-target, *t*(27) = 4.19, *p* < .001, *d* = 0.52 and new CRs, *t*(27) = 4.57, *p* < .001, *d* = 0.65, and the latter were statistically indistinguishable, *t*(27) = 0.95, *p* = .420, *d* = 0.13, suggesting that recollection was selective.

**Fig.2.**
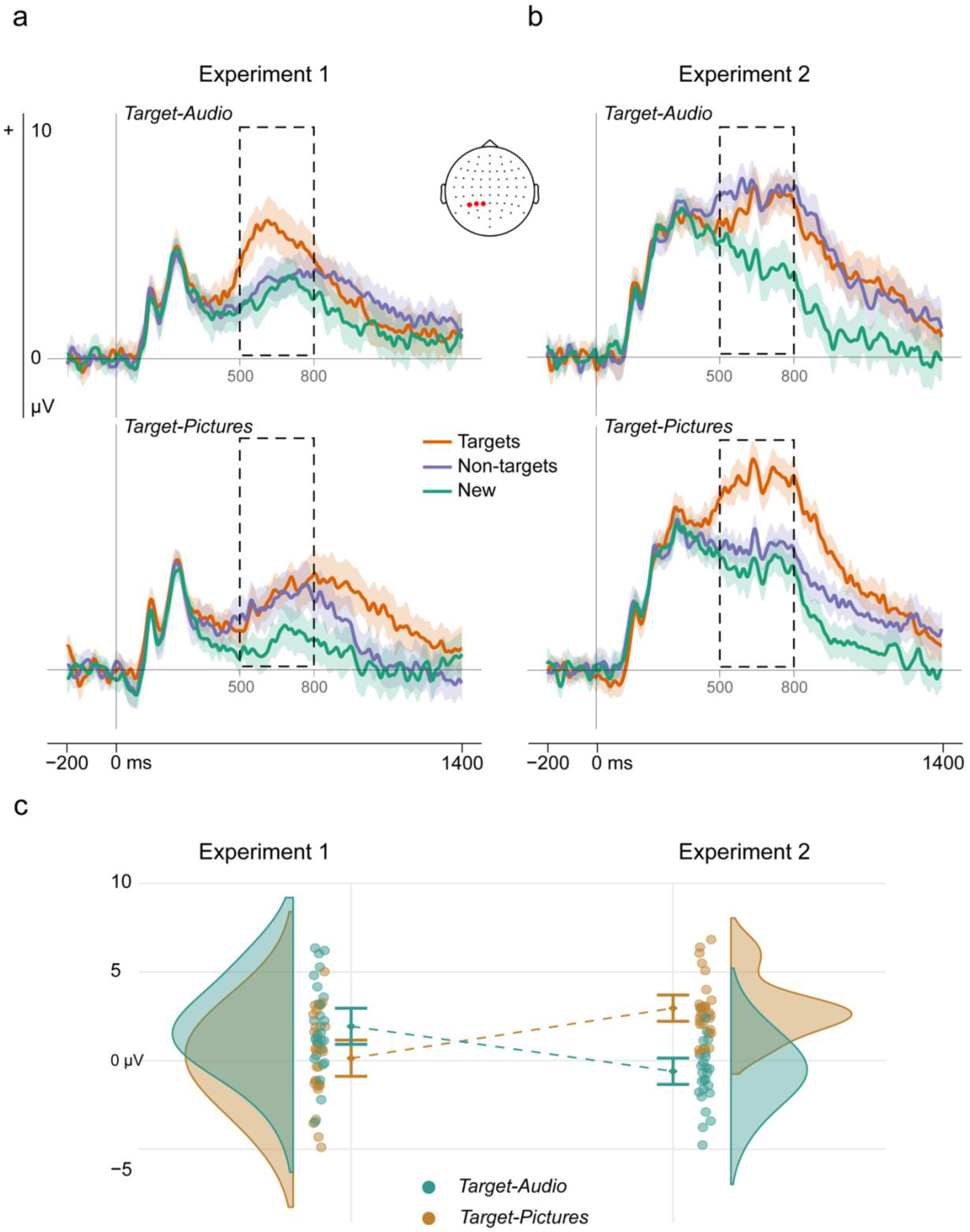
Selectivity of left parietal old/new effects. Panels (a) and (b) show mean grand-average ERP waveforms for target hits, non-target CRs, and new CRs over electrode sites (P1, P3, P5), plotted separately by Target Designation: (a) Experiment 1, with word cues, and (b) Experiment 2, with pictorial line drawing cues. The dashed areas indicate the analyzed time-window. The shaded areas represent the 95% confidence intervals for each time-point and adjusted using the Cousineau-Morey method for within-subject variables (*c*.*f*. Craddock, 2016). (c) Interaction effect of Target Designation x Cue Type (experiment) on the difference between left parietal ERPs evoked by target hits and non-target CRs. The colored dots are the difference scores for each participant, the shaded areas are the probability density function of the data, and error bars are the adjusted within-subject 95% confidence intervals around the means. Mean number of trials (range) contributing to ERPs in Experiment 1 for targets, non-targets, and new were 31 (14-37), 33 (18-39), and 35 (21-40) in the Target-Pictures block, and 29 (18-38), 34 (25-40), 33 (21-40) in the Target-Audio block. In Experiment 2, these were 33 (25-38), 32 (22-36), and 35 (25-40) in the Target-Pictures block, and 25 (15-34), 34 (30-38), 29 (16-38) in the Target-Audio block.

We interpreted the results of Experiment 1 in terms of the higher overlap between word cues and the auditory source, and so predicted a complementary asymmetry when retrieval cues were line drawings. The results of Experiment 2 supported this prediction (Fig. 2b,c). ANOVA with the same factors revealed a significant main effect of Item Type, *F*(1.55, 41.97) = 51.93, *p* < .001, *η*_*p*_^*2*^ = .658, a non-significant main effect of Target Designation *F*(1, 27) = 0.03, *p* = .874, *η*_*p*_^*2*^ = .001, and once again a significant interaction *F*(1.90, 51.21) = 20.98, *p* < .001, *η*_*p*_^*2*^ = .437. *Post hoc t-tests* showed that as expected, for the Target-Audio block ERPs to both target hits and non-target CRs were significantly more positive than new CRs, *t*(27) = 5.16, *p* < .001, *d* = 0.57, and *t*(27) = 5.60, *p* < .001, *d* = 0.67. Target ERPs were non-significantly more *negative*-going than non-target ERPs, *t*(27) = − 1.74, *p* = .094, *d* = 0.13, so recollection was nonselective. In contrast, target prioritization was significant, and more pronounced, in the Target-Pictures block. Here, ERPs evoked by target hits were significantly larger than those for non-target CRs, *t*(27) = 9.22, *p* < .001, *d* = 0.64, although both were significant relative to new CRs, *t*(27) = 9.25, *p* < .001, *d* = 0.84, and *t*(27) = 2.55, *p* = .018, *d* = 0.21.

These apparent differences from Experiment 1 were confirmed in a direct comparison (Fig. 2c). ANOVA with the additional between-participants factor of Cue Type (word cues/picture cues) on the difference between target and non-target ERPs revealed a significant interaction of Cue Type and Target Designation *F*(1, 54) = 38.04, *p* < .001, *η*_*p*_^*2*^ = .413. *Post hoc t-* tests confirmed that target and non-target left parietal effects differed more in the Target-Audio condition when cues were words (Experiment 1) as opposed to line drawings (Experiment 2), *t*(54) = 4.39, *p* < .001, *d* = 1.17, while selection in the Target-Picture condition was stronger when cues were line drawings, *t*(54) = − 5.01, *p* < .001, *d* = 1.34.

#### Global analysis

The results converged with the focal analyses to reveal complementary differences in target versus non-target old/new effects according to target designation in both experiments, as well as a further, later-onsetting difference in Experiment 1 (Fig. 3). When visual words were cues (Experiment 1), the difference between target and non-target ERPs was significantly greater when participants targeted auditory words than pictures (*p* = .004). This interaction cluster was present between 451-874 ms and was widespread across the scalp. Therefore, although cluster tests do not provide precise spatial or temporal localization (Sassenhagen & Draschkow, 2019), the effect overlapped that shown in our focal analyses. *Post hoc* tests within blocks revealed significantly larger target than non-target ERPs (*p* = .002) when participants targeted auditory words. There was also an unexpected later-onsetting significant interaction in the opposite direction in a cluster that was maximal over posterior electrodes and significant from 900-1,400 ms (*p* = .008). *Post hoc* tests revealed more positive-going target than non-target ERPs when participants targeted pictures (*p* = .020), but a non-significant reversed difference when participants targeted auditory words (*p* = .207).

**Fig. 3.**
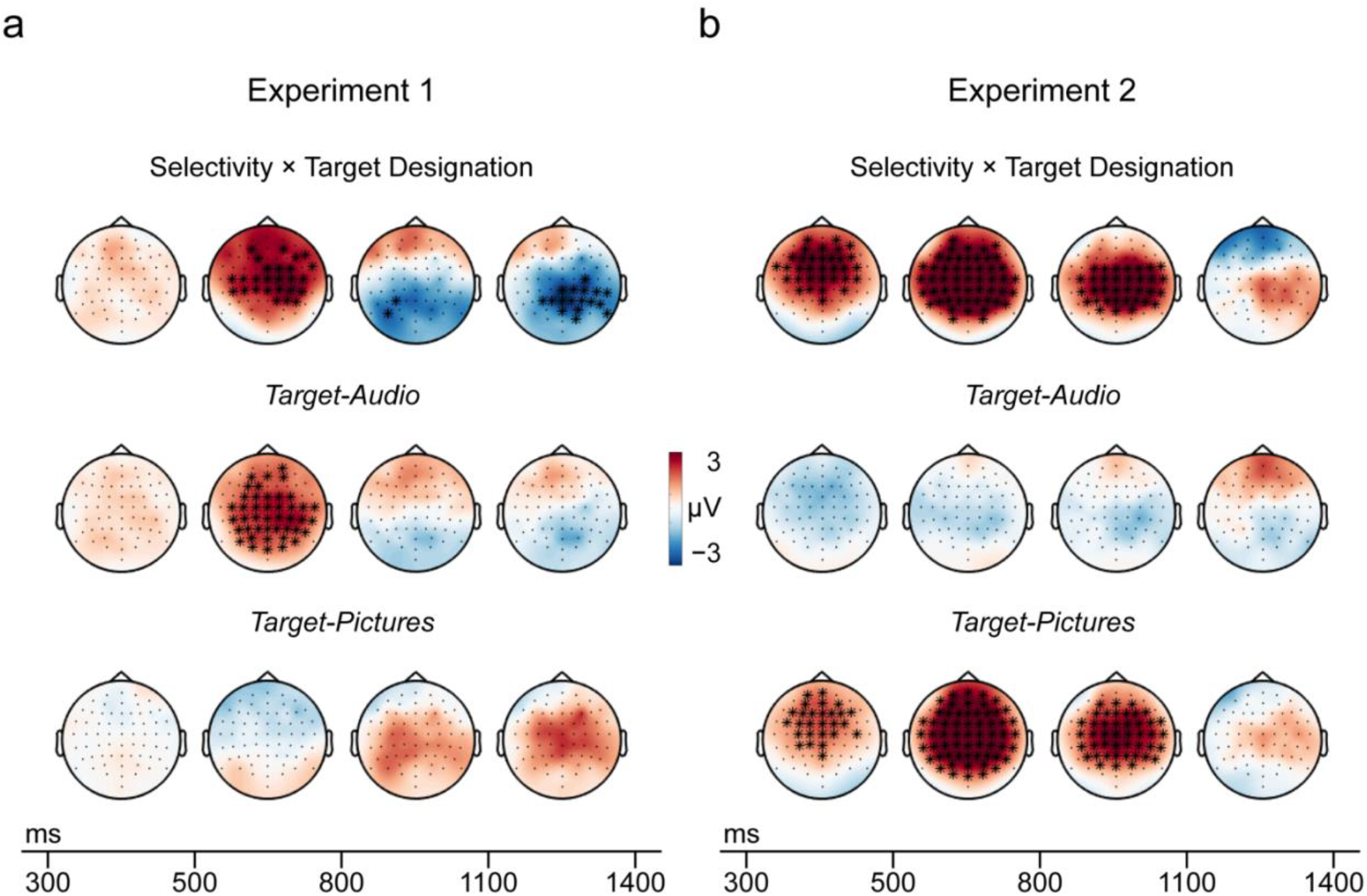
Global analyses of recollection selectivity: Target minus non-target ERPs. Topographic maps with significant clusters are shown for Experiment 1 (a) and Experiment 2 (b). Maps show ERP amplitude differences between target hits and non-target CRs by target designation (interaction at the top, Target-Audio in the middle, Target-Pictures at the bottom). Cluster significance is depicted by time-window, i.e., electrodes belonging to a significant cluster are highlighted (*) if significant on average in the plotted time-window.

In Experiment 2, as in the focal analysis, the difference between target and non-target ERPs was greater when pictures than auditory words were targets (*p* < .001). This effect was widespread over the scalp from 300-1,096 ms. *Post hoc* tests revealed that target and non-target ERPs did not differ reliably when participants used line drawings to target auditory words (*p* = .203), but when the same cues were used to target pictures, ERPs were more positive-going for targets (*p* < .001; Fig. 3b). This interaction cluster therefore overlapped the effect shown in the focal analysis and took the same form.

### Retrieval goal states

#### Focal analysis

Retrieval orientation ERP effects suggesting differences in retrieval goal states were present in both experiments and followed the predicted reversed pattern, although they were smaller and later-onsetting in Experiment 2 (Fig. 4). When visual words were cues (Experiment 1), ERPs to new CRs were more positive-going in the Target-Audio than the Target-Picture condition from about 400-1,300 ms with a centroparietal scalp maximum. When pictures were cues (Experiment 2), these ERPs were more positive-going in the Target-Picture than the Target-Audio condition from about 600-1,100 ms, with a frontal scalp maximum. Thus, ERPs to unstudied items were more positive-going in the high cue overlap condition in each experiment.

**Fig. 4.**
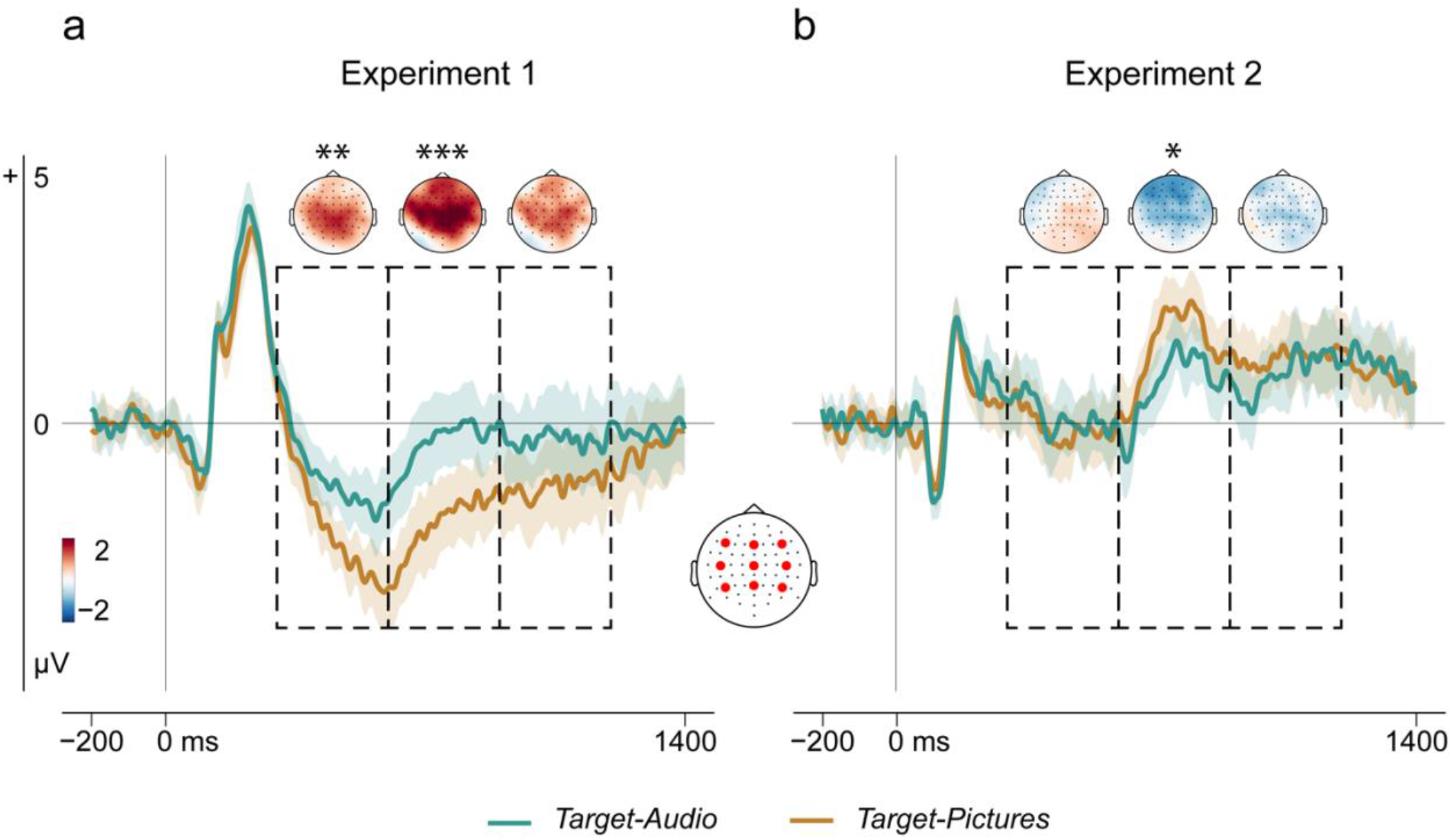
Retrieval goal states. Panels (a) and (b) show the results of the focal analysis. Grand-average ERP waveforms for retrieval orientation effects are plotted for (a) Experiment 1, word cues, and (b) Experiment 2, pictorial cues. Data are averaged over the grid of 9 electrodes, highlighted in red. The dashed areas indicate the time-windows from 300-600, 600-900 and 900-1200 ms, and time-windows with significant differences highlighted (* *p* ≤ .05, ** *p* ≤ .01, *** *p* ≤ .001). The shaded areas show 95% confidence intervals for each time-point and adjusted using the Cousineau-Morey method for within-subject variables (*c*.*f*. Craddock, 2016). The upper parts of each panel show topographic maps of ERP amplitude differences between new CRs according to target designation (Target-Audio minus Target-Pictures) in each experiment. + *p* ≤ .10, * *p* ≤ .05, ** *p* ≤ .01, *** *p* ≤ .001.

In Experiment 1, ANOVA with factors of Target Designation (picture/audio), Hemisphere (left/midline/right) and Site (anterior/central/posterior) showed a significant main effect of Target Designation in Experiment 1 between 300-600 ms, *F*(1,27) = 10.25, *p* = .003, *η*_*p*_^*2*^ = 275. This persisted from 600-900 ms, *F*(1, 27) = 12.76, *p* = .001, *η*_*p*_^*2*^ = .321, but was no longer reliable by 900-1200 ms post-stimulus, *F*(1,27) = 3.06, *p* = .091, η_p_^2^ = .102. In contrast, when cues were pictures (Experiment 2), retrieval orientation effects were significant only in the 600-900 ms time-window; for main effect of target designation, *F*(1, 27) = 4.39, *p* = .046, *η*_*p*_^*2*^ = .140; for 300-600 ms, *F*(1, 27) = 0.26, *p* = .613, *η*_*p*_^*2*^ = 0.01; for 900-1200 ms, *F*(1, 27) = 0.69, *p* = .413, *η*_*p*_^*2*^ = 0.025. Full ANOVA outputs are given in Supplemental Results available online (Table S1).

#### Global analysis

The results of the global retrieval orientation analyses also converged with those of the focal analyses. In Experiment 1, ERPs evoked by new CRs were more positive in the Target-Audio than the Target-Pictures block in a centroparietal cluster encompassing 403-920 ms (*p* = .002). However, for Experiment 2, the smaller retrieval orientation effects found in our focal analyses were not statistically significant in the global analysis (*p* = .064).

## Discussion

Recollecting the past involves selecting from a large number of stored memory traces. We used ERPs to investigate how and when people recover the desired information. By measuring time-resolved neural responses during recollection, we were able to quantify selection acting prior to this point. In two preregistered experiments, we manipulated the representational overlap between cues and the information to be remembered. In Experiment 1, visual word test cues shared more processing with studied auditory words than studied pictures. The left parietal effect, an ERP marker of recollection, was selective – larger for targets than non-targets – only when auditory words were targets, and not when pictures were targets. Importantly, this asymmetric pattern of selectivity could not be explained by easier recollection of targets, as regardless of target designation, responses were slightly more accurate and faster to items studied as pictures. The data thus favored a cue-overlap account, suggesting that at least with word cues, pre-retrieval selection was effective only for the high-overlap (auditory word) source. In the second experiment, we directly tested this interpretation by changing the cues at test to picture line drawings which would overlap more with the picture than the auditory source. As expected, we found target-selective left parietal effects only for the high-overlap (in this case picture) source.

These ERP asymmetries are consistent with previous studies showing selectivity of the left parietal effect for targets which were identical to “copy-cues” shown at test (Herron and Rugg 2003a; Stenberg et al., 2006). Together, the data support the view that cue-target overlap is a critical factor enabling selection prior to recollection. Moreover, we show for the first time that the degree of cue-target overlap has downstream consequences for successful retrieval even when overlap is incomplete (see also Czernochowski et al., 2005). These findings extend support for longstanding principles of memory which assume that cues trigger recollection when they elicit representations that overlap with stored memory traces (Morris, Bransford, & Franks, 1977; Tulving & Thomson, 1973). The principle of overlap is qualified by evidence that it is diagnostic rather than absolute overlap that must be maximized, so effective cues are also those that overlap more with the targeted *relative* to the non-targeted information (Goh & Lu, 2012; Nairne, 2002). In the current experiments, we increased diagnosticity by increasing cue-target overlap and reducing cue-non-target overlap in tandem, so did not test this further principle. This will be an important goal for future studies.

Theoretical models of recollection assume that selection is achieved via goal-directed elaboration on available external cues (Anderson & Bjork, 1994). Both the current experiments showed differences in ERPs according to retrieval orientation – the current retrieval goal – on new item trials where no retrieval took place. ERPs were more positive-going in the conditions with the greatest cue-target overlap, extending previous findings from studies with one copy-cue condition (Herron & Rugg, 2003a; Hornberger et al., 2004; see also Morcom & Rugg, 2012). Dzulkifli and Wilding (2005, 2006) previously demonstrated retrieval orientation ERP effects only when recollection was selective, and abolished both these goal-related effects and selective recollection by increasing target retrieval difficulty. However, our data suggest that the difficulty account is insufficient. The cue overlap account could explain Dzulkifli and Wilding’s (2005, 2006) findings if the ‘difficulty’ manipulations prevented proactive goal-directed processes from generating effective cues for one source, for example because longer study lists make source cues less diagnostic. This proposal remains to be tested. A related question concerns the nature of the goal-driven processing participants engaged to maximize diagnostic overlap. Participants may have elaborated external cues by mentally reinstating representations stored in the targeted memory, for example by emphasizing the phonological features shared between word cues and auditorily studied items in Experiment 1. Alternatively, or in addition, they may have constrained cue processing to *decrease* overlap with non-targeted memory representations, for example by suppressing imagery processes that would overlap with non-targeted picture representations in Experiment 1 (Hornberger et al., 2004). The present data cannot distinguish between these two accounts.

Although our data demonstrate clearly complementary patterns of selective recollection of the two sources under different retrieval cues, questions also remain about the degree of selection achieved. While in Experiment 1, recollection as measured using the left parietal ERP effect appeared to be completely selective for auditory word targets, in Experiment 2 the same marker suggested incomplete selectivity in the corresponding high-overlap picture target condition (Fig. 2). There is no reason to think that selective remembering is an all-or-none phenomenon, and the magnitude of the left parietal ERP effect has been shown to track the amount (Vilberg & Rugg, 2009) and precision (Murray et al., 2015) of recollected information. Although prior ERP studies that used visual word copy-cues to target studied visual words (Herron and Rugg, 2003a; Stenberg et al., 2006) did not detect a non-target left parietal ERP effect when non-targets were pictures, an fMRI study using the same task revealed a more nuanced picture. Although only targets elicited old/new effects in left angular gyrus (a possible source of the left parietal ERP effect), non-target old/new effects were detected in other brain regions (Morcom & Rugg, 2012). Experiment 1’s findings echo this earlier result but add additional temporal information: although the left parietal effect was selective in the target-audio condition with no detectable effect for non-targets, the global analysis showed that there was some processing of non-targets after 800 ms post-stimulus. This later-onsetting activity presumably reflected post-retrieval processing. Other studies have also reported incomplete selectivity of the left parietal effect (see Rosburg and Mecklinger, 2017). Further research is needed to determine whether some elements of memories are more easily selected than others, and to understand potential trade-offs between proactive pre-retrieval processing and reactive post-retrieval processing that may have consequences for populations who are less able to remember selectively (see Morcom, 2016).

In conclusion, these experiments are the first to show that recollection is selective when retrieval cues overlap more closely with sought-for information in memory, implicating pre-retrieval control. When recollection is selective, neural activity associated with retrieval goal states is also more pronounced. The data open up several new possibilities for future research into the goal-states that enable selective remembering, its implementation, and consequences for mnemonic experience.

## Supplemental Information

Supplemental materials and results are available online and can be accessed at https://osf.io/8pt25/.

## Acknowledgments

The authors would like to thank Gintare Siugzdinyte for her contribution to the initial design of the second experiment.

## Open practices statement

The design, analysis plan and hypotheses for the experiments were pre-registered on the Open Science Framework (OSF) and can be accessed at https://osf.io/j84z6 (Experiment 1) and https://osf.io/pqn4z (Experiment 2). The stimuli and task materials as well as data and analysis scripts were also posted on the OSF and will be made publicly available upon publication in a scientific journal.

## Declarations

### Ethics approval

All procedures performed in these studies were in line with the principles of the Declaration of Helsinki. Approval was granted by the Psychology Research Ethics Committee at the University of Edinburgh, ref.: 135-1819/1 (Experiment 1) and 300-1819/1 (Experiment 2).

### Consent to participate

Informed consent was obtained from all individual participants included in the studies.

### Consent for publication

The authors confirmed that participants provided informed consent for publication of their anonymized data.

### Availability of data and materials

The stimuli, task materials and data are publicly available and can be accessed online.

### Code availability

Task and analysis scripts are publicly available and can be accessed online.

### Authors’ contribution

A. Moccia and A.M. Morcom developed the study concept and design. Testing and data collection were performed by A. Moccia. A. Moccia performed the data analysis under the supervision of A.M. Morcom. A. Moccia drafted the manuscript and A.M. Morcom and A. Moccia provided critical revisions. Both authors approved the final version of the manuscript for submission.

## Supplemental Materials

### Supplemental image processing

Line drawings were created by first applying a 5 Hz high-pass filter to the images in Photoshop. Filtered images were then transformed to black and white drawings, and the white background was set as the alpha channel so that the images appeared as black line drawings on a transparent background. Image processing on filtered pictures was achieved in Matlab 2018a. In Experiment 1, 76 pictures were taken from the BOSS database (Brodeur et al., 2014), 68 from the POPORO database (Kovalenko et al., 2012), and 96 were sourced online. In Experiment 2, seven practice items were swapped to the main experimental task to improve image quality of test cues. Of these, four additional images were taken the POPORO database (Kovalenko et al., 2012) and three were sourced online.

### Supplemental EEG pre-processing

#### Thresholds for automated artifact rejection

The pre-registered EEG preprocessing pipeline and artefact rejection criteria can be found at osf.io/j84z6 and osf.io/pqn4z (see EEG recording and pre-processing for more details). We used customized functions from the FASTER toolbox (Nolan et al., 2010) to identify and reject epochs with excessive motion artefacts, drifts, or gross EOG movements, and channels with excessive noise. For each participant, we rejected epochs whose *z*-transformed values exceeded ± 3 over trials. These values were each epoch’s amplitude range, variance, and deviation from each channel’s mean value. The latter measure was computed by subtracting each channel’s mean amplitude within the epoch from the channel’s mean amplitude across epochs, which was then averaged across all channels. Channels’ thresholds were computed by calculating each channel’s mean correlation with all other channels, variance, and Hurst exponent, a measure of a signal’s long-range dependence.

## Supplemental Results

### Discrimination performance in the Recognition Exclusion Task

In a separate preregistered analysis of memory performance using discrimination (d’) measures to correct for response bias, we calculated *d’* for item memory, source memory, and nontarget false recognition for the Target-Picture and Target-Audio blocks (Snodgrass & Corwin, 1988). Item d’ was calculated by subtracting the z-scored proportion of new FAs from the *z*-scored proportion of target hits. Similarly, source d’ was computed by subtracting the *z*-nontargets FAs score from the *z*-target hits score, and nontarget d’ was obtained by subtracting the *z*-new FAs score to *z*-nontarget FAs score. Before computing these *d’* measures, raw trial numbers were corrected for a potential outcome of zero by adding 1 to the sum of old and the sum of new items and 0.5 to the target Hits, nontarget CRs, or nontarget FAs (Hautus, 1995; Snodgrass & Corwin, 1988). Confidence intervals (CIs) reported below are adjusted using the Cousineau-Morey method for within-subject variables (Morey, 2008).

The results of the *d*’ analysis converged with those of the analysis of response proportions reported in the main paper (see Exclusion Task Performance). We examined whether participants’ mnemonic discrimination scores differed according to target designation with paired-sample *t*-tests. Memory was consistently better for pictures compared to auditory words. In Experiment 1, while participant’s source memory was not significantly different between the targeted pictures (*M* = 2.14, 95% CI = 0.24) and targeted auditory words (*M* = 2.19, 95% CI = 0.21), *t*(27) = − 0.39, *p* = .700, their item memory was higher for pictures (*M* = 2.40, 95% CI = 0.19) than auditory words (*M* = 2.05, 95% CI = 0.22), *t*(27) = 2.16, *p* = .040, despite the greater overlap of test cues with the auditory source condition. There was also a significant increase in false recognition of nontargets when these had been studied as auditory words (*M* = 0.26, 95% CI = 0.28) as opposed to pictures (*M* = − 0.14, 95% CI = 0.26), *t*(27) = 2.98, *p* = .006.

In Experiment 2, there was again consistently better memory for pictures than auditory words. Here, both item memory and source memory were significantly higher when participants targeted pictures (item memory: *M* = 2.65, 95% CI = 0.18, source memory: *M* = 2.46, 95% CI = 0.18) versus auditory words (item memory: *M* = 1.58, 95% CI = 0.18, source memory: *M* = 2.19, 95% CI = 0.21), *t*(27) = 7.71, *p* < .001 and *t*(27) = 2.37, *p* = 0.025, respectively. This increased mnemonic discrimination accuracy for pictures than auditory words was consistent with the greater overlap of test cues with the picture source. Once again, false recognition was greater for items that that been studied as auditory words (*M* = 0.19, 95% CI = 0.20) as opposed to pictures (*M* = − 0.60, 95% CI = 0.25), *t*(27) = 5.67, *p* < .001.

### Retrieval goal states supplemental results

#### Full ANOVA results of focal analysis

To investigate retrieval orientation effects indexing retrieval goal states, we analyzed ERPs elicited by new CRs according to target designation in a focal analysis (see ERP results in main manuscript for details). Full ANOVA outputs are given in Table S1.

**Table S1.** Focal Retrieval Orientation Analyses: Omnibus Repeated Measures ANOVAs

| Time<br>window | Effect | Experiment 1 |  |  |  |  | Experiment 2 |  |  |  |  |
| --- | --- | --- | --- | --- | --- | --- | --- | --- | --- | --- | --- |
| | | df | MSE | *F* | $\eta p2$ | *p* | df | MSE | *F* | $\eta p2$ | *p* |
| 300 - 600 |  |  |  |  |  |  |  |  |  |  |  |
|  | Target Designation | 1, 27 | 12.30 | 10.25<br>** | .275 | .003 | 1, 27 | 19.38 | 0.26 | .010 | .613 |
|  | Hemisphere | 1.93,<br>52.00 | 9.72 | 5.89<br>** | .179 | .005 | 1.97,<br>53.19 | 13.83 | 1.66 | .058 | .200 |
|  | Site | 1.11,<br>30.09 | 52.03 | 40.88<br>*** | .602 | <.001 | 1.08,<br>29.12 | 75.20 | 90.66<br>*** | .771 | <.001 |
|  | Target<br>Designation×Hemisphere | 1.97,<br>53.30 | 1.42 | 0.43 | .016 | .652 | 1.88,<br>50.75 | 1.28 | 3.18<br>+ | .105 | .053 |
|  | Target Designation×Site | 1.22,<br>32.81 | 3.12 | 1.75 | .061 | .196 | 1.34,<br>36.17 | 1.93 | 1.63 | .057 | .212 |
|  | Hemisphere×Site | 2.85,<br>76.83 | 1.86 | 31.11<br>*** | .535 | <.001 | 3.16,<br>85.39 | 3.24 | 7.17<br>*** | .210 | <.001 |
|  | Target<br>Designation×Hemisphere×Site | 3.47,<br>93.62 | 0.27 | 0.85 | .030 | .485 | 3.13,<br>84.59 | 0.58 | 0.27 | .010 | .858 |
| 600 - 900 |  |  |  |  |  |  |  |  |  |  |  |
|  | Target Designation | 1, 27 | 27.81 | 12.76<br>** | .321 | .001 | 1, 27 | 17.16 | 4.39<br>* | .140 | .046 |
|  | Hemisphere | 1.99,<br>53.85 | 15.30 | 4.04<br>* | .130 | .023 | 1.95,<br>52.68 | 13.77 | 0.87 | .031 | .425 |
|  | Site | 1.21,<br>32.59 | 30.10 | 60.96<br>*** | .693 | <.001 | 1.07,<br>28.90 | 62.18 | 18.07<br>*** | .401 | <.001 |
|  | Target<br>Designation×Hemisphere | 1.84,<br>49.56 | 2.42 | 0.12 | .004 | .872 | 1.99,<br>53.66 | 1.59 | 0.33 | .012 | .717 |
|  | Target Designation×Site | 1.27,<br>34.34 | 3.64 | 1.36 | .048 | .260 | 1.31,<br>35.49 | 3.57 | 1.13 | .040 | .312 |
|  | Hemisphere×Site | 2.23,<br>60.09 | 4.11 | 29.32<br>*** | .521 | <.001 | 3.12,<br>84.21 | 4.24 | 30.44<br>*** | .530 | <.001 |
|  | Target<br>Designation×Hemisphere×Site | 3.62,<br>97.63 | 0.50 | 0.75 | .027 | .547 | 3.31,<br>89.37 | 0.66 | 0.58 | .021 | .646 |
| 900 - 1,200 |  |  |  |  |  |  |  |  |  |  |  |
|  | Target Designation | 1, 27 | 37.53 | 3.06<br>+ | .102 | .092 | 1, 27 | 21.04 | 0.69 | .025 | .413 |
|  | Hemisphere | 1.95,<br>52.56 | 13.73 | 10.42<br>*** | .278 | <.001 | 1.87,<br>50.57 | 14.30 | 0.95 | .034 | .388 |
|  | Site | 1.25,<br>33.64 | 18.44 | 28.16<br>*** | .511 | <.001 | 1.08,<br>29.16 | 41.41 | 4.85<br>* | .152 | .033 |
|  | Target<br>Designation×Hemisphere | 1.94,<br>52.46 | 1.94 | 0.19 | .007 | .819 | 1.94,<br>52.47 | 2.33 | 0.06 | .002 | .943 |
|  | Target Designation×Site | 1.65,<br>44.63 | 2.50 | 1.38 | .049 | .260 | 1.43,<br>38.58 | 3.93 | 0.07 | .003 | .874 |
|  | Hemisphere×Site | 2.72,<br>73.54 | 3.42 | 18.83<br>*** | .411 | <.001 | 3.46,<br>93.45 | 3.35 | 31.02<br>*** | .535 | <.001 |
|  | Target<br>Designation×Hemisphere×Site | 3.57,<br>96.51 | 0.62 | 1.24 | .044 | .299 | 3.70,<br>99.97 | 0.79 | 1.32 | .046 | .271 |

+ $p \leq .10$ , \* $p \leq .05$ , \*\* $p \leq .01$ , \*\*\* $p \leq .001$ .

### Preparatory cue ERP analyses

Further pre-registered analyses of retrieval orientation effects examined differential preparatory activity when participants were preparing to retrieve according to target designation (i.e, when pictures were targets versus when auditory words were targets, see Materials and Procedure in main manuscript). These ERPs were time-locked to pre-cues that were followed by a correct response to the subsequent trial in the Target-Picture and Target-Audio block. The mean number of trials and range that contributed to these ERPs were 98.46 (65-112) and 96.43 (69-113) for Target-Picture and Target-Audio pre-cues, respectively, in Experiment 1 and 102.31 (87-115) and 91.66 (75-106) in Experiment 2. In a focal analysis, we examined ERPs over a grid of 6 frontal electrodes (F1, F3, F5/F2, F4, F6) in four 250 ms time-windows from 0-1,000 ms, following Herron (2018). ANOVAs with factors of Target Designation (picture/audio) and Hemisphere (left/right) did not yield any significant results including the factor of Target Designation (for full results see Table S2). In the global analysis across all scalp electrodes and time-points in the preparatory time-window (0-2,300 ms), we used a cluster-based permutation *t*-test to correct for multiple comparisons (Maris & Oostenveld, 2007; Oostenveld et al., 2011, see Global ERP analyses in main manuscript). These analyses also did not show any significant effects of Target Designation (at 1-tailed cluster-level *α* of .025).

**Table S2.** Focal Repeated Measures ANOVAs: Preparatory Cue Effects

| Time window (ms) | Effect | Experiment 1 |  |  |  |  | Experiment 2 |  |  |  |  |
| --- | --- | --- | --- | --- | --- | --- | --- | --- | --- | --- | --- |
| | | df | MSE | *F* | $\eta^2_{\text{partial}}$ | *p* | df | MSE | *F* | $\eta^2_{\text{partial}}$ | *p* |
| 0 - 250 | Target Designation | 1, 27 | 1.04 | 0.97 | .035 | .333 | 1, 27 | 0.91 | 0.14 | .005 | .712 |
|  | Hemisphere | 1, 27 | 0.14 | 8.22 ** | .233 | .008 | 1, 27 | 0.08 | 1.38 | .049 | .251 |
|  | Target Designation×Hemisphere | 1, 27 | 0.05 | 0.01 | <.001 | .913 | 1, 27 | 0.06 | 0.97 | .035 | .335 |
| 250 - 500 | Target Designation | 1, 27 | 1.93 | 0.01 | <.001 | .907 | 1, 27 | 1.93 | 1.29 | .046 | .266 |
|  | Hemisphere | 1, 27 | 0.22 | 19.85 *** | .424 | <.001 | 1, 27 | 0.39 | 1.91 | .066 | .178 |
|  | Target Designation×Hemisphere | 1, 27 | 0.16 | 0.23 | .008 | .636 | 1, 27 | 0.14 | 0.49 | .018 | .491 |
| 500 - 750 | Target Designation | 1, 27 | 1.86 | 0.12 | .004 | .737 | 1, 27 | 1.44 | 0.30 | .011 | .590 |
|  | Hemisphere | 1, 27 | 0.29 | 25.51 *** | .486 | <.001 | 1, 27 | 0.26 | 8.82 ** | .246 | .006 |
|  | Target Designation×Hemisphere | 1, 27 | 0.15 | 0.00 | <.001 | .971 | 1, 27 | 0.18 | 1.08 | .038 | .308 |
| 750 - 1,000 | Target Designation | 1, 27 | 2.17 | 0.01 | <.001 | .907 | 1, 27 | 1.86 | 2.08 | .071 | .161 |
|  | Hemisphere | 1, 27 | 0.39 | 17.88 *** | .398 | <.001 | 1, 27 | 0.23 | 10.42 ** | .278 | .003 |
|  | Target Designation×Hemisphere | 1, 27 | 0.14 | 0.04 | .001 | .851 | 1, 27 | 0.23 | 0.73 | .026 | .400 |

+ $p \leq .10$ , \* $p \leq .05$ , \*\* $p \leq .01$ , \*\*\* $p \leq .001$ .

## References

Anderson, M.C., Bjork, R.A., 1994. Mechanisms of inhibition in long term memory: A New Taxoomy. In: Dagenbach, D., Carr, T.H. (Eds.), Inhibitory Processes in Attention, Memory, and Language. Academic Press, San Diego, pp. 265–325.

Craddock, M., (2016, November 28). ERP Visualization: Within-subject confidence intervals. https://www.mattcraddock.com/blog/2016/11/28/erp-visualization-within-subject-confidence-intervals/

Benjamini, Y., & Hochberg, Y. (1995). Controlling the False Discovery Rate: A Practical and Powerful Approach to Multiple Testing. Journal of the Royal Statistical Society: Series B (Methodological), 57(1), 289–300. https://doi.org/10.1111/j.2517-6161.1995.tb02031.x

Brodeur, M. B., Guérard, K., & Bouras, M. (2014). Bank of Standardized Stimuli (BOSS) Phase II: 930 New Normative Photos. PLoS ONE, 9(9), e106953. https://doi.org/10.1371/journal.pone.0106953

Craik, F. I. M. (1983). On the Transfer of Information from Temporary to Permanent Memory [and Discussion]. Philosophical Transactions of the Royal Society of London, 302(1110), 341–359.

Czernochowski, D., Mecklinger, A., Johansson, M., & Brinkmann, M. (2005). Age-related differences in familiarity and recollection: ERP evidence from a recognition memory study in children and young adults. Cognitive, Affective, & Behavioral Neuroscience, 5(4), 417–433. https://doi.org/10.3758/CABN.5.4.417

Delorme, A., & Makeig, S. (2004). EEGLAB: an open source toolbox for analysis of single-trial EEG dynamics including independent component analysis. In Journal of Neuroscience Methods (Vol. 134).

Dunlap, W. P., Cortina, J. M., Vaslow, J. B., & Burke, M. J. (1996). Meta-Analysis of Experiments With Matched Groups or Repeated Measures Designs. Psychological Methods, 1(2), 170–177.

Dywan, J., Segalowitz, S. J., & Webster, L. (1998). Source Monitoring: ERP Evidence for Greater Reactivity to Nontarget Information in Older Adults. Brain and Cognition, 36(3), 390–430. https://doi.org/10.1006/brcg.1997.0979

Dzulkifli, M. A., Herron, J. E., & Wilding, E. L. (2006). Memory retrieval processing: Neural indices of processes supporting episodic retrieval. Neuropsychologia, 44(7), 1120–1130. https://doi.org/10.1016/j.neuropsychologia.2005.10.021

Dzulkifli, M. A., & Wilding, E. L. (2005). Electrophysiological indices of strategic episodic retrieval processing. Neuropsychologia, 43(8), 1152–1162. https://doi.org/10.1016/j.neuropsychologia.2004.11.019

Evans, L. H., Wilding, E. L., Hibbs, C. S., & Herron, J. E. (2010). An electrophysiological study of boundary conditions for control of recollection in the exclusion task. Brain Research, 1324, 43–53. https://doi.org/10.1016/j.brainres.2010.02.010

Goh, W. D., & Lu, S. H. X. (2012). Testing the myth of the encoding–retrieval match. Memory & Cognition, 40(1), 28–39. https://doi.org/10.3758/s13421-011-0133-9

Herron, J. E., & Wilding, E. L. (2005). An Electrophysiological Investigation of Factors Facilitating Strategic Recollection. Journal of Cognitive Neuroscience, 17(5), 777–787. https://doi.org/10.1162/0898929053747649

Herron, Jane E., & Rugg, M. D. (2003a). Retrieval Orientation and the Control of Recollection. Journal of Cognitive Neuroscience, 15(6), 843–854. https://doi.org/10.1162/089892903322370762

Herron, Jane E., & Rugg, M. D. (2003b). Strategic influences on recollection in the exclusion task: Electrophysiological evidence. Psychonomic Bulletin & Review, 10(3), 703–710. https://doi.org/10.3758/BF03196535

Hornberger, M., Morcom, A. M., & Rugg, M. D. (2004). Neural Correlates of Retrieval Orientation: Effects of Study–Test Similarity. Journal of Cognitive Neuroscience, 16(7), 1196–1210. https://doi.org/10.1162/0898929041920450

Jacoby, L. L. (1991). A process dissociation framework: Separating automatic from intentional uses of memory. Journal of Memory and Language, 30(5), 513–541. https://doi.org/10.1016/0749-596X(91)90025-F

Keating, J., Affleck-Brodie, C., Wiegand, R., & Morcom, A. M. (2017). Aging, working memory capacity and the proactive control of recollection: An event-related potential study. PLOS ONE, 12(7), e0180367. https://doi.org/10.1371/journal.pone.0180367

Kovalenko, L. Y., Chaumon, M., & Busch, N. A. (2012). A Pool of Pairs of Related Objects (POPORO) for Investigating Visual Semantic Integration: Behavioral and Electrophysiological Validation. Brain Topography, 25(3), 272–284. https://doi.org/10.1007/s10548-011-0216-8

Maris, E., & Oostenveld, R. (2007). Nonparametric statistical testing of EEG- and MEG-data. Journal of Neuroscience Methods, 164(1), 177–190. https://doi.org/10.1016/j.jneumeth.2007.03.024

Morcom, A. M. (2016). Mind Over Memory: Cuing the Aging Brain. Current Directions in Psychological Science, 25(3), 143–150. https://doi.org/10.1177/0963721416645536

Morcom, A. M., & Rugg, M. D. (2012). Retrieval Orientation and the Control of Recollection: An fMRI Study. Journal of Cognitive Neuroscience, 24(12), 2372–2384. https://doi.org/10.1162/jocn_a_00299

Morey, R. D. (2008). Confidence Intervals from Normalized Data: A correction to Cousineau (2005). Tutorials in Quantitative Methods for Psychology, 4(2), 61–64. https://doi.org/10.20982/tqmp.04.2.p061

Morris, C. D., Bransford, J. D., & Franks, J. J. (1977). Levels of processing versus transfer appropriate processing. Journal of Verbal Learning and Verbal Behavior, 16(5), 519–533. https://doi.org/10.1016/S0022-5371(77)80016-9

Murray, J. G., Howie, C. A., & Donaldson, D. I. (2015). The neural mechanism underlying recollection is sensitive to the quality of episodic memory: Event related potentials reveal a some-or-none threshold. NeuroImage, 120, 298–308. https://doi.org/10.1016/j.neuroimage.2015.06.069

Nairne, J. S. (2002). The myth of the encoding-retrieval match. Memory, 10(5–6), 389–395. https://doi.org/10.1080/09658210244000216

Nolan, H., Whelan, R., & Reilly, R. B. (2010). FASTER: Fully Automated Statistical Thresholding for EEG artifact Rejection. Journal of Neuroscience Methods, 192(1), 152– 162. https://doi.org/10.1016/j.jneumeth.2010.07.015

Nuwer, M. R., Comi, G., Emerson, R., Fuglsang-Frederiksen, A., Guérit, J.-M., Hinrichs, H., Ikeda, A., Jose C. Luccas, F., & Rappelsburger, P. (1998). IFCN standards for digital recording of clinical EEG. Electroencephalography and Clinical Neurophysiology, 106(3), 259–261. https://doi.org/10.1016/S0013-4694(97)00106-5

Oostenveld, R., Fries, P., Maris, E., & Schoffelen, J.-M. (2011). FieldTrip: Open Source Software for Advanced Analysis of MEG, EEG, and Invasive Electrophysiological Data. Computational Intelligence and Neuroscience, 2011. https://doi.org/10.1155/2011/156869

R Core Team (2019). R: A language and environment for statistical computing. R Foundation for Statistical Computing, Vienna, Austria. URL http://www.R-project.org/

Rosburg, T., & Mecklinger, A. (2017). The costs of target prioritization and the external requirements for using a recall-to-reject strategy in memory exclusion tasks: A meta-analysis. Psychonomic Bulletin and Review, 24(6), 1844–1855. https://doi.org/10.3758/s13423-017-1256-1

Rosburg, T., Mecklinger, A., & Johansson, M. (2011). Strategic retrieval in a reality monitoring task. Neuropsychologia, 49(10), 2957–2969. https://doi.org/10.1016/j.neuropsychologia.2011.07.002

Rugg, M. D., & Curran, T. (2007). Event-related potentials and recognition memory. Trends in Cognitive Sciences, 11(6), 251–257. https://doi.org/10.1016/j.tics.2007.04.004

Rugg, M. D., & Wilding, E. L. (2000). Retrieval processing and episodic memory. Trends in Cognitive Sciences, 4(3), 108–115. https://doi.org/10.1016/S1364-6613(00)01445-5

Sassenhagen, J., & Draschkow, D. (2019). Cluster-based permutation tests of MEG/EEG data do not establish significance of effect latency or location. Psychophysiology, 56(6), e13335. https://doi.org/10.1111/psyp.13335

Sprondel, V., Kipp, K. H., & Mecklinger, A. (2012). Electrophysiological evidence for late maturation of strategic episodic retrieval processes: Late maturation of strategic retrieval. Developmental Science, 15(3), 330–344. https://doi.org/10.1111/j.1467-7687.2011.01130.x

Stenberg, G., Johansson, M., & Rosén, I. (2006). Conceptual and perceptual memory: Retrieval orientations reflected in event-related potentials. Acta Psychologica, 122(2), 174–205. https://doi.org/10.1016/j.actpsy.2005.11.001

Tulving, E., & Thomson, D. M. (1973). Encoding specificity and retrieval processes in episodic memory. Psychological Review, 80(5), 352–373. https://doi.org/10.1037/h0020071

Unsworth, N. (2016). The Many Facets of Individual Differences in Working Memory Capacity. In Psychology of Learning and Motivation (Vol. 65, pp. 1–46). Elsevier. https://doi.org/10.1016/bs.plm.2016.03.001

Vega-Mendoza, M., West, H., Sorace, A., & Bak, T. H. (2015). The impact of late, non-balanced bilingualism on cognitive performance. Cognition, 137, 40–46. https://doi.org/10.1016/j.cognition.2014.12.008

Vilberg, K. L., & Rugg, M. D. (2009). Functional significance of retrieval-related activity in lateral parietal cortex: Evidence from fMRI and ERPs. Human Brain Mapping, 30(5), 1490–1501. https://doi.org/10.1002/hbm.20618

Wilding, E. L., Fraser, C. S., & Herron, J. E. (2005). Indexing strategic retrieval of colour information with event-related potentials. Cognitive Brain Research, 25(1), 19–32. https://doi.org/10.1016/j.cogbrainres.2005.04.012

## References

Hautus, M. J. (1995). Corrections for extreme proportions and their biasing effects on estimated values ofd′. Behavior Research Methods, Instruments, & Computers, 27(1), 46–51. https://doi.org/10.3758/BF03203619

Herron, J. E. (2018). Direct electrophysiological evidence for the maintenance of retrieval orientations and the role of cognitive control. NeuroImage, 172, 228–238. https://doi.org/10.1016/j.neuroimage.2018.01.062

Snodgrass, J. G., & Corwin, J. (1988). Pragmatics of Measuring Recognition Memory: Applications to Dementia and Amnesia. Journal of Experimental Psychology: General, 117(1), 34–50.

